# cGAS deficient mice display premature aging associated with de-repression of LINE1 elements and inflammation

**DOI:** 10.1101/2024.10.10.617645

**Authors:** John C. Martinez, Francesco Morandini, Lucinda Fitzgibbons, Natasha Sieczkiewicz, Sung Jae Bae, Michael E. Meadow, Eric Hillpot, Joseph Cutting, Victoria Paige, Seyed Ali Biashad, Matthew Simon, John Sedivy, Andrei Seluanov, Vera Gorbunova

**Affiliations:** Translational Biomedical Sciences Program, University of Rochester, NY, 14627, USA; Department of Biology, University of Rochester, NY, 14627, USA; Department of Molecular Biology, Cell Biology, and Biochemistry, Brown University, RI, 02912, USA; Department of Medicine, University of Rochester, NY, 14627, USA

## Abstract

Aging-associated inflammation, or ‘inflammaging” is a driver of multiple age-associated diseases. Cyclic GMP-AMP Synthase (cGAS) is a cytosolic DNA sensor that functions to activate interferon response upon detecting viral DNA in the cytoplasm. cGAS contributes to inflammaging by responding to endogenous signals such as damaged DNA or LINE1 (L1) cDNA which forms in aged cells. While cGAS knockout mice are viable their aging has not been examined. Unexpectedly, we found that cGAS knockout mice exhibit accelerated aging phenotype associated with induction of inflammation. Transcription of L1 elements was increased in both cGAS knockout mice and in cGAS siRNA knockdown cells associated with high levels of cytoplasmic L1 DNA and expression of ORF1 protein. Cells from cGAS knockout mice showed increased chromatin accessibility and decreased DNA methylation on L1 transposons. Stimulated emission depletion microscopy (STED) showed that cGAS forms nuclear condensates that co-localize with H3K9me3 heterochromatin marks, and H3K9me3 pattern is disrupted in cGAS knockout cells. Taken together these results suggest a previously undescribed role for cGAS in maintaining heterochromatin on transposable elements. We propose that loss of cGAS leads to loss of chromatin organization, de-repression of transposable elements and induction of inflammation resulting in accelerated aging.

## Introduction

Properly regulating inflammatory gene expression is critical to defending the organism’s overall health and strength. During aging, chronic overproduction of pro-inflammatory cytokines and genes such as IL-6, IL-8, IL-13, type-one interferons (IFNs), and NF-kB among others results in a debilitating phenotype known as ‘inflammaging,’ one of the major hallmarks of aging ^1,2^. As the proportion of the population over 65 years old continues to increase at a rapid rate, inflammaging will pose an increasing burden to the healthcare system and quality of life^3^.

Cyclic GMP-AMP Synthase (cGAS) is a cytosolic sensor that regulates cellular response to double-stranded nucleic acids in the cytoplasm^4^. Upon binding nucleic acids in the cytoplasm, cGAS produces cyclic GMP-AMP (cGAMP) which then binds to Stimulator of Interferon Genes (STING) on the endoplasmic reticulum (ER)^5^. Following cGAMP binding, STING becomes translocated from the ER to the Golgi Apparatus where it facilitates phosphorylation of IRF3 via TBK1. This phosphorylation induces IRF3 nuclear import and subsequent transcription of multiple inflammatory genes, including NF- kB and IL-6^6^. While cGAS is intended to target exogenous DNA, the Mab21 domain of cGAS (responsible for binding nucleic acids and generating cGAMP) is unable to decern between viral and endogenous nucleic acids, including double-stranded L1 transcripts in the cytoplasm^7^. Further, cGAS has also been shown to activate upon binding mitochondrial DNA, which is known to occur during cellular aging and as a driver of neuroinflammation^8,9^. Having demonstrated its implication in the immune response stemming from L1 cDNA, cGAS has also emerged as a potential pharmacological candidate for auto-immune diseases including systematic lupus erythematosus (SLE) and psoriasis^10, 11^. Due to these unintended, chronic inflammatory effects, both cGAS and STING have been proposed as targets to manage healthy aging, autoimmune disease, and prevention of certain cancers^12,13^.

Long interspersed nuclear element 1 (LINE1 or L1) is a class 1 transposable element that comprises around 20% of both the mouse and human genomes^14, 15^. While only a minority of full-length (around 6 kb) elements in mice and humans are capable of fully independent retrotransposition, L1s persist largely as thousands of partially truncated elements scattered throughout the genome. Full-length L1s encode for two proteins, ORF1p, a nucleic acid chaperone, and ORF2p, an endonuclease and reverse transcriptase (RT)^16,17,18,19^. When fully functional, translation of these elements results in the formation of a ribonucleoprotein (RNP) comprising both ORF proteins and the L1 mRNA that encoded them. This RNP complex is then imported back into the nucleus, where ORF2p creates a single strand break before utilizing its RT capabilities to copy L1 cDNA back into the genome from the 3’ end. Because of epigenomic regulation, relatively few full-length L1 elements, and difficulties transcribing both strands of the L1 sequence back into the genome, a successful retrotransposition event is exceedingly rare relative to L1 mRNA transcription.

Cells have also evolved to be able to co-opt the unavoidable presence of L1 elements in the genome, which has led to their presence being conceptualized as a ‘double-edged sword^20^. Although the blind mole rat, a long-lived rodent, has been shown to utilize L1 transcriptional activation as an anti-cancer mechanism, L1 activation has also been demonstrated to be a significant driver of inflammation in SIRT6 deficient mice, which exhibit an accelerated aging phenotype defined by a severe lifespan decrease and growth defect, as well as during cellular senescence, another hallmark of aging^7, 21^. L1 activation has also been shown to be necessary for proper embryonic development and serves as an essential mediator of neuronal plasticity^22,23^. Despite potential benefits of L1 activity, cells must maintain tight control over L1s to avoid cellular damage. Accordingly, cells have evolved numerous mechanisms to repress L1 expression via proteins important in genome maintenance and tumor suppression, including SIRT6, KAP1, and p53^24, 25^. In the gametes specifically, cells express PIWI factors, a class of proteins responsible for additional repression of L1 expression by acting as a guide for siRNAs, piRNAs, and miRNAs targeting L1 mRNA for degradation^26^. Highlighting the importance of proper regulation of L1s and other transposable elements, cells have evolved hundreds of KRAB zinc-finger proteins to systematically repress the promotors of these elements^27^. This widespread, systematic repression of L1 expression in both somatic and germ cells demonstrate the importance of proper L1 regulation.

Given the potential effects cGAS may have on longevity and healthspan, we sought to explore the relationship between cGAS knockout, cellular inflammatory signaling, and inflammaging. Specifically, we aimed to better understand the relationship between cGAS deficiency, L1 activity, and inflammation on cellular aging and maintenance. Surprisingly, our findings show that knockdown or knockout of cGAS, both *in vitro* and *in vivo*, results in aberrant L1 transcription and cytosolic cDNA accumulation. Additionally, RNAseq and ATACseq data from cGAS KO, but not STING KO, mice reveal a dramatic increase in the accessibility of pro-inflammatory genes and L1 elements, suggesting a previously unknown role for chromatin-bound cGAS in regulating inflammatory response. qRT-PCR confirmed the increase in pro-inflammatory genes in cGAS KO, but not STING KO, mice. The unexpected upregulation of L1 transcripts upon cGAS deletion subsequently results in chromatin disorganization, activation of L1 elements, DNA damage, and induction pro-inflammatory signaling. Moreover, nuclear based cGAS appears to undergo liquid-liquid phase separation (LLPS) in the nucleus, and these structures dissociate upon treatment with 1,6 hexanediol, commonly used to disrupt phase separation in cells^28^. Taken together, our data uncovers that nuclear-based cGAS plays a critical role in suppressing L1 transcription and maintaining chromatin organization.

## Results

### cGAS KO mice exhibit accelerated-aging phenotype and increased inflammation

The effect of long-term cGAS deletion on mouse health has not been sufficiently characterized. Apart from increased susceptibility to viral infections, cGAS KO mice have not been reported to display any physiological or morphological deficiencies compared to their wild-type counterparts^29^. As downregulation of cGAS-STING pathway may be a strategy to counteract inflammaging we set out to characterize the effects of cGAS KO on murine healthspan.

Middle aged (18 months old) cGAS KO mice of both sexes displayed significant increase in frailty index compared to wild type mice, with males exhibiting a higher degree of frailty compared to females (**Fig. 1a, Sup. Fig. 1a, b**)^30^. This increase in frailty in cGAS KO mice was driven by an increased incidence of kyphosis, piloerection, and tremors (**Sup. Fig. 1d**). Moreover, cGAS KO males displayed significant body weight increase and obesity (**Fig. 1b, f, Sup. Fig. 1c, f**). Finally, we conducted histological analysis of various organs from cGAS KO and WT mice. Compared to WT mice, lungs of cGAS KO mice exhibited elevated neutrophil infiltration within their inflamed tissue (**Fig. 1c**). Elevated inflammation and tissue fibrosis was also observed in the liver (**Fig. 1d**). Finally, female cGAS KO mice displayed shortened lifespan (**Fig. 1f**). While male cGAS KO mice experienced increased inflammation, obesity, and frailty, their lifespan was not different from the WT (**Sup. Fig. 1e**). However, regardless of sex, cGAS KO mice experience severe detrimental outcomes over time. Collectively, these results demonstrate that cGAS KO mice display accelerated-aging phenotype characterized by increased frailty in both sexes and shortened lifespan in females.

**Figure 1.**
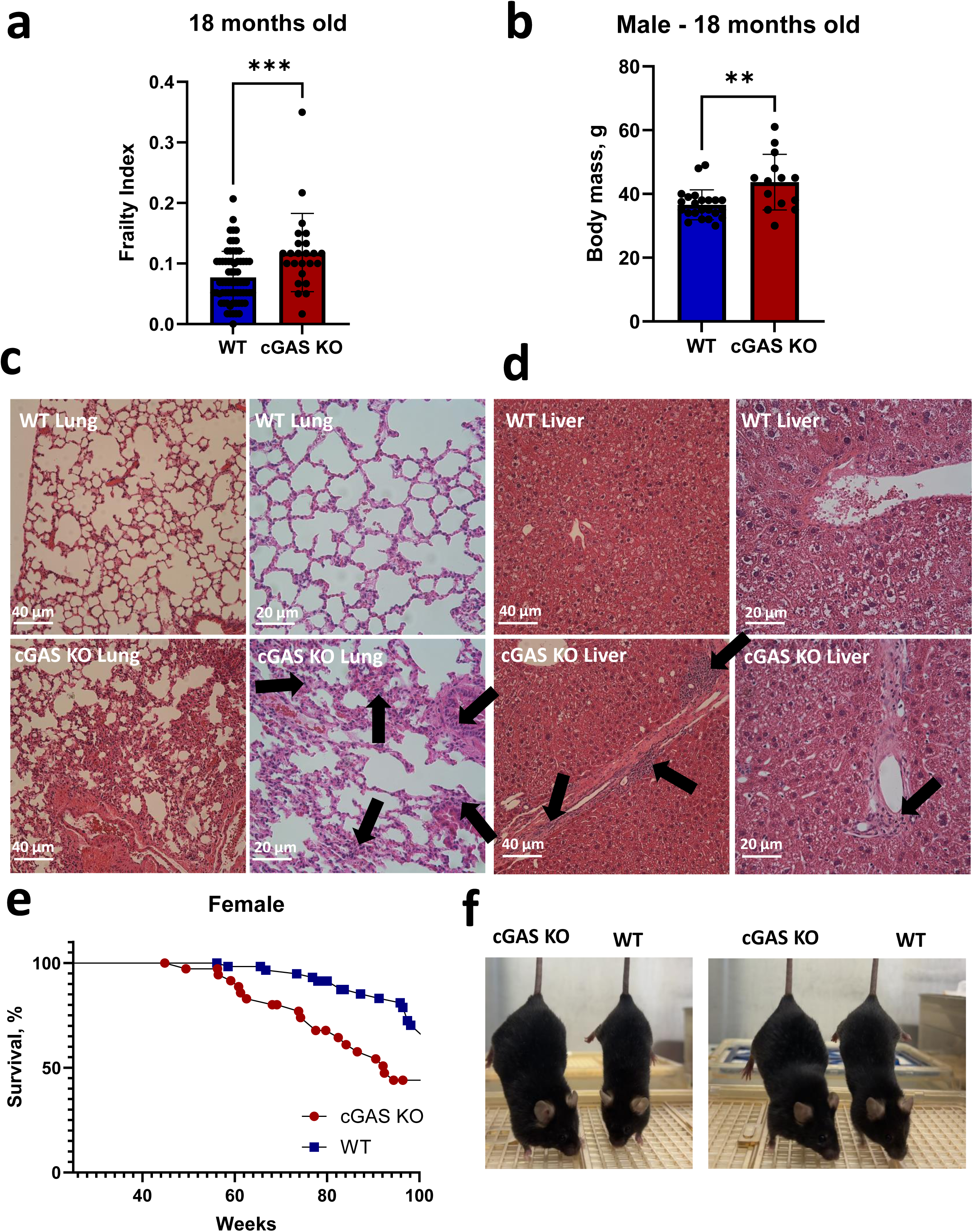
cGAS knockout mice show increased frailty and inflammation. (a) Frailty index in 18-months old WT and cGAS KO mice. cGAS KO mice display increased frailty. p < 0.001, t-test. WT n = 64 mice; cGAS KO n = 24 mice. (b) cGAS KO male mice display higher body mass at older age. p < 0.01, t-test. WT n = 23 mice; cGAS KO n =14 mice. (c) H&E staining of lung in WT and cGAS KO mice. Relative to WT, cGAS KO tissues display substantial neutrophil infiltration (indicated by arrows). (d) H&E staining of liver in WT and cGAS KO mice. Relative to WT, cGAS KO livers display substantial increase in fibrotic tissue (indicated by arrows). (e) Survival of WT and cGAS KO female mice. cGAS KO mice show a significantly reduced lifespan (WT n = 48, cGAS KO n = 58; WT median survival = 106.7 weeks, cGAS KO survival = 92.4 weeks; p = 0.0008, Mantel-Cox test). (f) Representative images of two different pairs of cGAS KO and WT male mice at 14-months-old.

### RNAseq reveals pro-inflammatory signature in cGAS KO mice

To further characterize the effect of cGAS knockout we performed RNA sequencing on primary lung cells derived from 12-month-old mice (**Sup. Fig. 2a; Source Data 1**). We chose to work on isolated fibroblasts to focus on cell-intrinsic effects while avoiding effects due to the differences in immune infiltration we showed in **Figure 1c**. To ensure the composition of these cells reflected their organ of origin as closely as possible, cells were isolated from their tissues immediately following the sacrifice of the animal, cultured in DMEM F12, and passaged once before analysis to ensure removal of residual tissue material from the samples. We found 315 upregulated and 190 downregulated genes in cGAS KO compared to WT (Adjusted p-value < 0.05, |Log2FC| > 0.5, **Fig. 2a; Source Data 2**). Gene set enrichment analysis (GSEA) showed upregulation of biological processes related to inflammation (**Fig. 2b; Source Data 3**). Downregulated biological processes, on the other hand, were related to cytoskeletal organization. We noticed that upregulated genes were enriched for components of the inflammasome complex (Aim2, Nlrp3, Nlrp1a/b, Casp1, etc. **Fig. 2c; Source Data 4**). The inflammasome is a large protein complex that induces inflammation, mainly through activation of IL-1b and IL-18, upon stimulation by pathogens and damage associated molecular patterns (PAMPs, DAMPs)^31^. Aim2 in particular is an inflammasome subunit that senses cytosolic double-stranded DNA, similar to cGAS. We confirmed by qRT-PCR that Aim2 and downstream targets (IL-18, NFkB, TNFα) were also upregulated in lung tissue (**Fig. 2d**). We suspect that upregulation of Aim2 could be compensating for the loss of cGAS. Prior studies discovered crosstalk between the Aim2 inflammasome and the cGAS-STING pathway, and similar to our observations, Aim2 was found to be elevated in a lupus mouse model upon cGAS knockout^32,10,33,34^. However, it remains unclear why Aim2 would not only be upregulated, but also activated.

**Figure 2.**
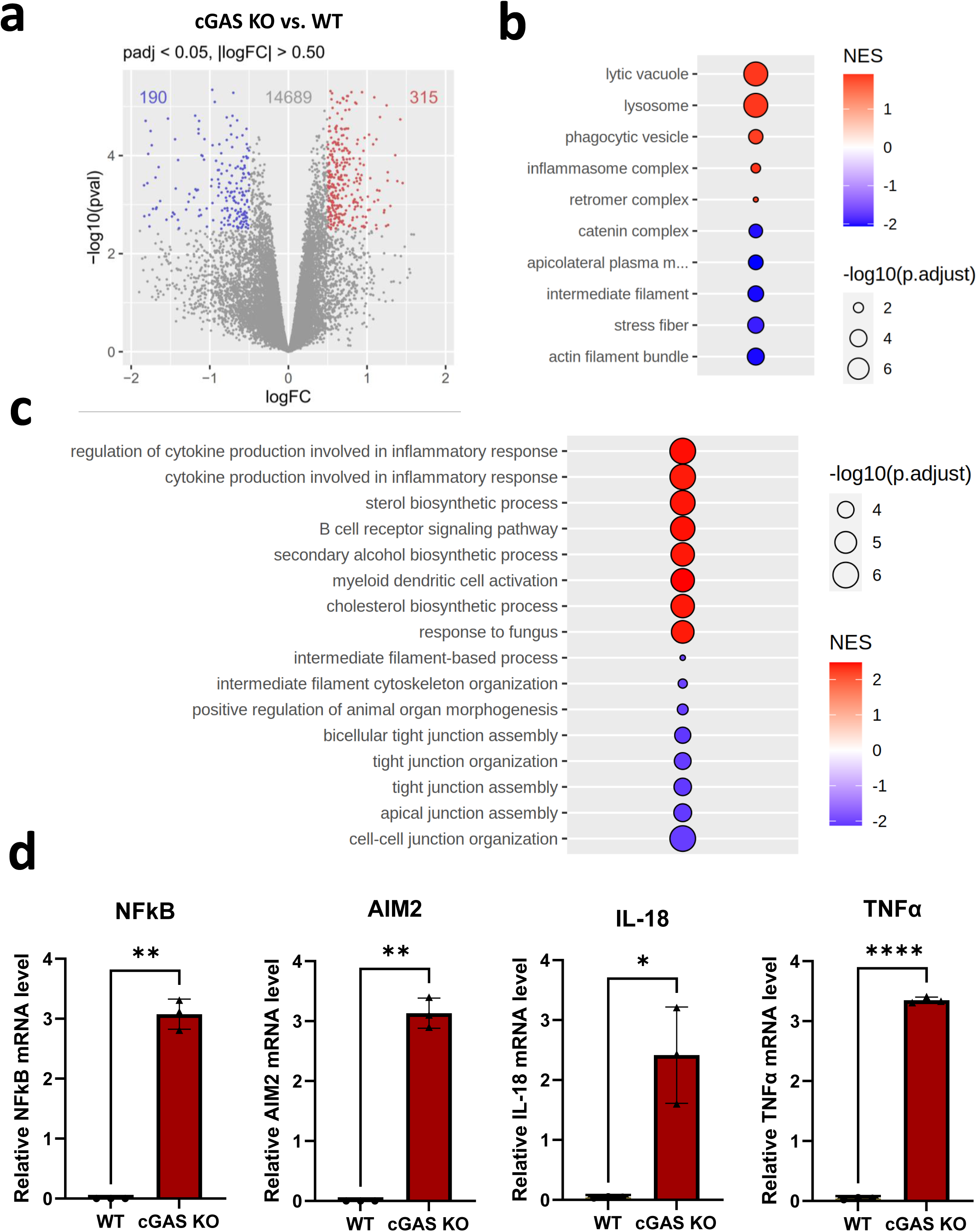
RNAseq reveals pro-inflammatory signature in cGAS KO mice. (a) Volcano plot showing differentially expressed genes in lung cells from cGAS KO vs. WT mice. Relative to WT, cGAS KO samples display 315 significantly upregulated genes and 190 downregulated genes. cGAS KO n = 6 mice; WT n = 5 mice. (b) Top biological processes GO Terms for cGAS KO versus WT mouse lung cells show pro-inflammatory signature in cGAS KO samples. (c) Top cellular components GO Terms for cGAS KO versus WT mouse lung cells. (d) qRT-PCR confirms higher levels of inflammation-related gene expression in cGAS KO mice. NFkB, IL-18, and IL-6, along with the PYHIN gene AIM2, are significantly enriched in cGAS KO mice but not WT mice. qRT-PCR was conducted on 12-month-old WT and cGAS KO lung tissue. N=3 mice per group. *p < 0.05, **p < 0.01, or ****p < 0.0001, t-test.

We additionally asked if the inflammatory response we observed was coherent with the senescence induced secretory phenotype (SASP). To do so, we performed GSEA on a gene set characteristic of senescence^35^. Interestingly, senescence-associated genes were overall downregulated although not significantly (**Sup. Fig. 2b**). Thus, a senescent phenotype is likely not driving the inflammation experienced in cGAS KO cells, coherently with reports that cGAS is necessary to initiate senescence^36^.

### cGAS KO mice display increased LINE1 (L1) expression across multiple tissues

We hypothesized that L1s may be the endogenous source of inflammation as they have been reported to stimulate cytosolic DNA sensing pathways through accumulation of L1 cDNA in the cytoplasm. We quantified the expression of mouse L1 families in our fibroblast RNA-seq data and found upregulation of L1MdA_I, L1MdFanc_I and II (**Fig. 3a; Source Data 5**). L1MdA_I in particular is one of the most active L1 families in mice, and has pathogenic potential^37^. Quantitative qRT-PCR on lung, kidney and liver tissue of 12-month-old mice confirmed significant upregulation of L1MdA mRNA expression in cGAS KO mice (**Fig. 3b**). cGAS KO was verified via Western blot on lung tissue (**Sup. Fig. 3**).

**Figure 3.**
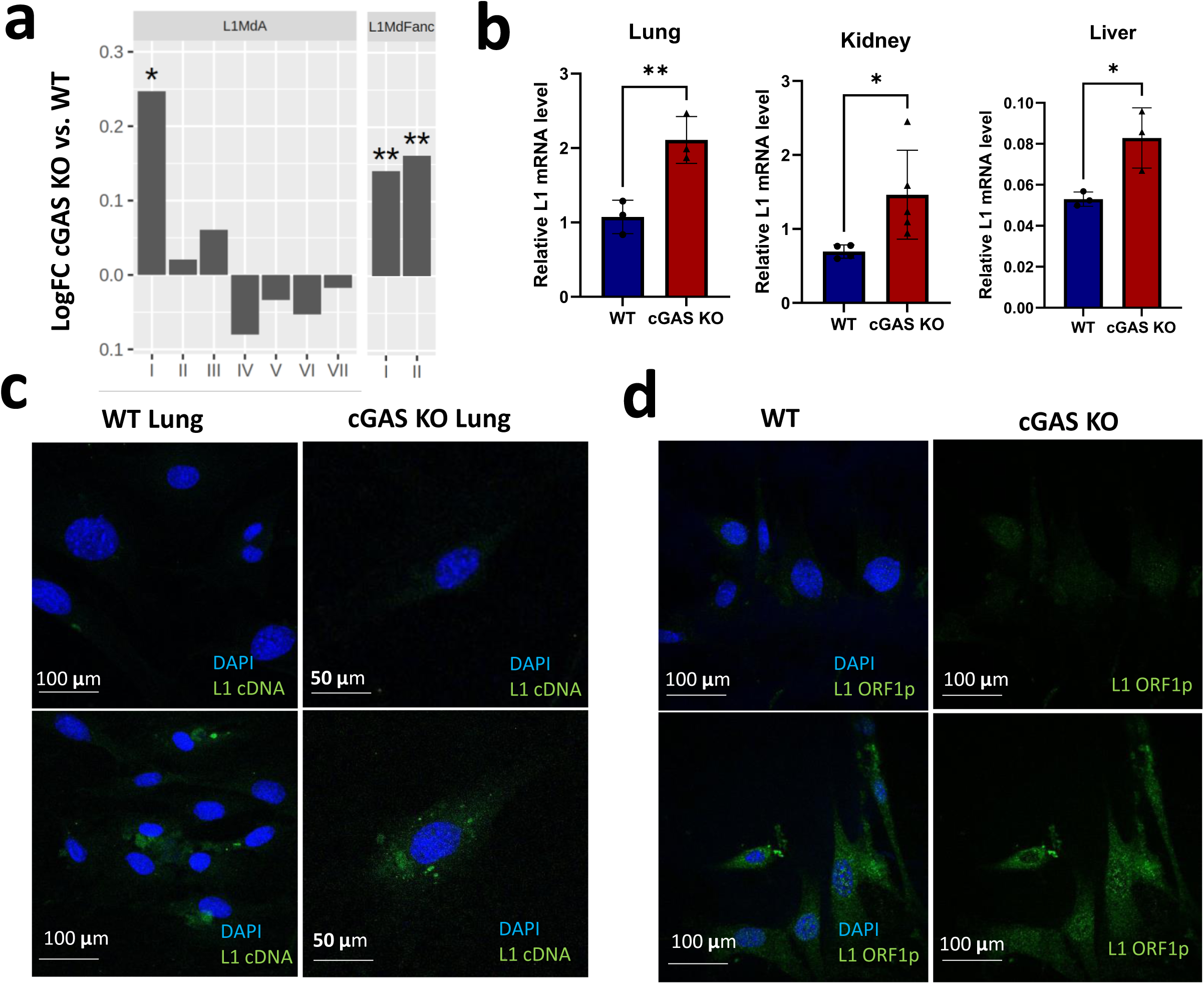
cGAS knockout mice display increased L1 expression across multiple tissues. (a) RNAseq analysis of primary lung cells from cGAS KO and WT mice shows changes in L1Md L1 families upon cGAS KO. L1 mRNA expression in L1MdA_I, L1Mda_II, L1MdA_III, and L1MdFanc families is higher in cGAS KO mouse lung. (b) qRT-PCR confirms elevated L1MdA expression in cGAS KO mouse tissues. Organs were harvested from 12-month-old WT and cGAS KO mice. L1 expression was measured via qRT-PCR and normalized to EEF2. n=3-6 mice per group. *p < 0.05 or **p < 0.01, t-test. (c) cGAS KO primary cells display elevated levels of L1 cDNA in the cytoplasm. smiFISH probes specific for L1 ORF1p and ORF2p L1MdA sequence were applied to WT and cGAS KO primary lung cells. Signal for cytoplasmic L1 cDNA was consistently elevated in cGAS KO cells. (d) Immunostaining of WT and cGAS KO mouse primary cells for ORF1p. Cells were stained using LINE1 ORF1p (EA13) antibody.

As L1 triggers inflammation via formation of cytoplasmic L1 cDNA we next tested whether the increased L1 mRNA expression led to cytoplasmic L1 cDNA formation.^7,38^ We conducted DNA smiFISH on WT and cGAS KO primary cells. smiFISH probes were designed to hybridize with L1 cDNA, and not hybridize to L1 mRNA transcripts to enhance specificity of the signal. Strikingly, smiFISH showed that cGAS KO primary cells display severely elevated L1MdA cDNA in the cytoplasm (**Fig. 3c**). We next tested if cGAS KO cells display increased ORF1p via IF. Consistent with smiFISH, cGAS KO cells showed increased ORF1p accumulation (**Fig. 3d**). These data demonstrate that cGAS KO mice display increased L1 transcription, cytoplasmic L1 cDNA accumulation, and ORF1p accumulation.

To confirm that L1 de-repression is a direct consequence of cGAS depletion we utilized siRNA to knock down cGAS in wild type mouse embryonic fibroblasts (MEFs) (**Fig. 4a**). We observed a nearly 4-fold increase in relative L1MdA mRNA expression (**Fig. 4b**). L1 cDNA smiFISH showed that knockdown of cGAS also resulted in L1MdA cDNA formation in the cytoplasm (**Fig. 4c, d**). The increase in cytoplasmic L1MdA cDNA and mRNA transcription was additionally complemented by a significant increase in ORF1p protein levels in cGAS knock-down MEFs (**Fig. 4e, f**).

**Figure 4.**
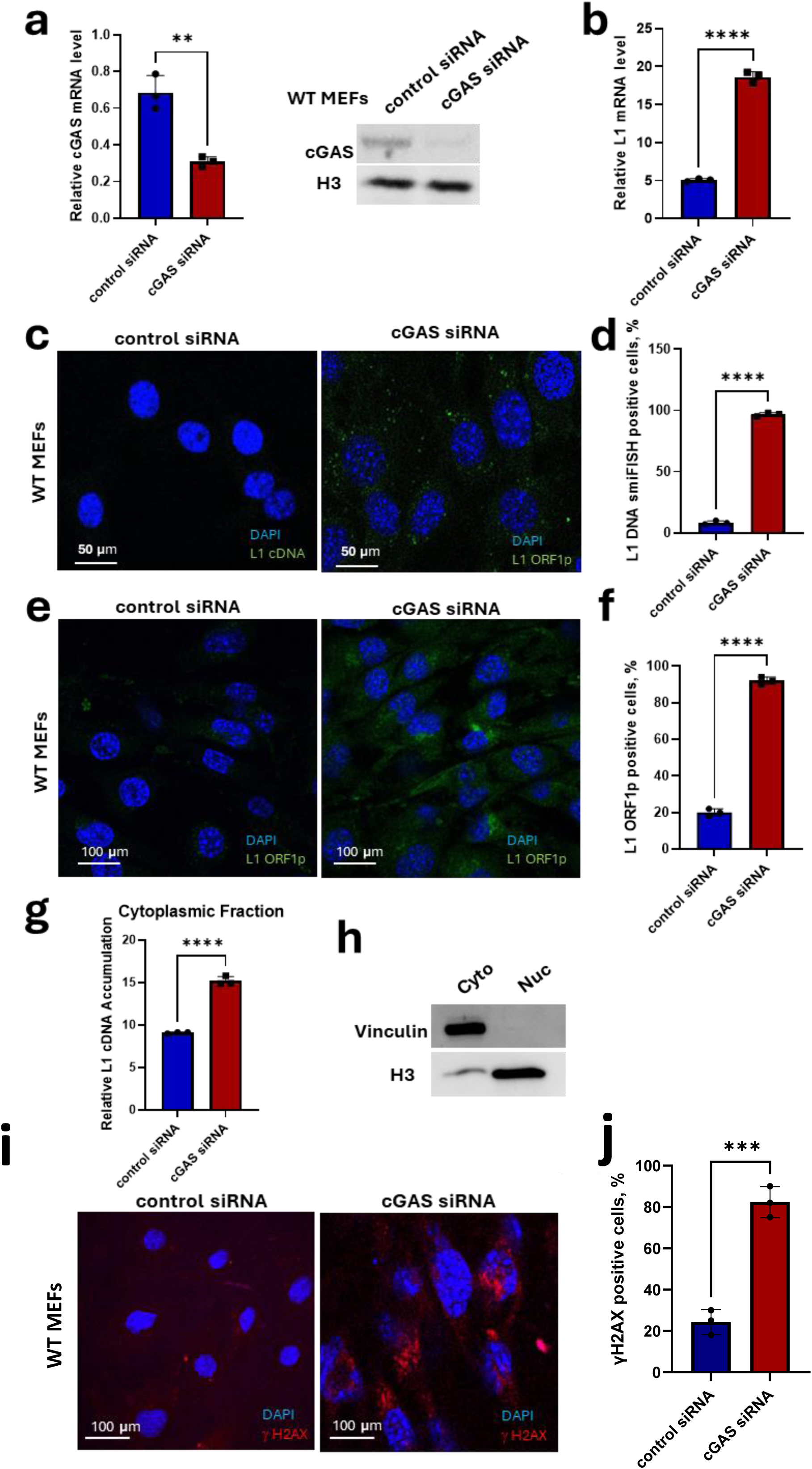
cGAS knockdown leads to aberrant L1 expression, accumulation of L1 cDNA, and DNA damage. (a) cGAS knockdown in MEFs. WT MEFs were treated with cGAS or control shRNA. cGAS knockdown was confirmed via qRT-PCR and Western blotting. (b) L1MdA mRNA expression was measured via qRT-PCR. EEF2 was used as a reference. (c) cGAS knockdown results in L1 cDNA accumulation. WT MEFs were treated with cGAS or control shRNA before conducting DNA smiFISH for L1MdA cDNA. Cells treated with cGAS shRNA display substantial increase in L1 cytoplasmic cDNA. (d) Quantification of (c). (e) cGAS knockdown results in L1 ORF1p accumulation. Immunostaining of WT MEFs treated with control or cGAS shRNA and stained for ORF1p (EA13). (f) Quantification of (e). (g) DNA qPCR following subcellular fractionation shows increased L1MdA cDNA in cytoplasm of cells treated with cGAS shRNA. MEFs were fractionated and cytoplasmic fraction was subjected to qPCR following DNA cleanup (Thermo) and RNase treatment. (h) Western blot to confirm successful cellular fractionation. Cytoplasmic and nuclear fractions were stained for vinculin and H3. (i) cGAS knockdown cells induces DNA damage signaling. gH2AX immunostaining of WT and cGAS KO MEFs transfected with control or cGAS shRNA. (j) Quantification of (i). Experiments were performed in triplicate **p < 0.01. ***p < 0.001, ****p < 0.0001, t-test

Additionally, we probed the presence of L1MdA cytoplasmic cDNAs via sub-cellular fractionation, DNA isolation and purification, followed by qPCR (**Fig. 4g**). Successful fractionation was verified via Western blotting (**Fig. 4h**). The cytosolic fraction of cells treated with cGAS siRNA demonstrated elevated levels of L1MdA cDNA relative to untreated MEFs. Taken together, the depletion of cGAS in cultured cells induced L1 de-repression similar to the one observed in cGAS KO mice. This result suggests a potential role chromatin-bound cGAS plays in regulating L1 repression. Finally, we observed accumulation of γH2AX in cells deficient for cGAS suggesting that the loss of cGAS results in DNA damage (**Fig. 4i, j**).

### Depletion of other cytoplasmic DNA sensors does not lead to L1 activation

After confirming that cGAS KO mice display elevated L1 levels, we set out to determine if deletion of other cytoplasmic DNA sensors has a similar effect. STING mediates the next step in the cGAS/STING inflammatory pathway, where it undergoes dimerization following cGAS activation upon binding nucleic acids in the cytoplasm^39^. The PYHIN family of genes, represents a separate class of cytosolic-DNA sensors that also signal via STING^40,41^. Remarkably, we found that both STING KO and PYHIN KO mice maintain repression of L1MdA family (**Sup. Fig. 4)**. This suggests that L1MdA de-repression is specifically associated with cGAS depletion, rather than a consequence of the overall cGAS/STING pathway disruption or disruption of other cytosolic DNA sensing proteins.

### Loss of cGAS results in major changes in chromatin accessibility

While cGAS plays essential role in cytoplasmic DNA sensing, the majority of cellular cGAS resides in the nucleus and is chromatin-bound^42^. Recent evidence has linked nuclear-based cGAS to DNA repair^43^. However, the effect of chromatin-bound cGAS on the epigenome has not been described.

To determine the effect of losing nuclear cGAS on chromatin accessibility, we performed ATACseq on cGAS KO and wild-type lung primary fibroblasts (**Source Data 6**). We detected 185219 open chromatin regions (OCRs), of which 1392 were more open in cGAS KO and 200 were less open in cGAS KO (**Fig. 5a; Source Data 7**). We performed gene set overrepresentation analysis based on genes downstream of opening promoters (OCRs within 1000 bp of gene transcription start sites) and found upregulation of inflammatory pathways (**Fig. 5b; Source Data 8**), in accordance with out RNA-seq results (**Fig. 2b**). For example, Aim2 gene encoding a cytoplasmic DNA sensor that stimulates expression of interferon genes, became more accessible in cGAS KO cells (**Fig. 5d**). No terms were enriched based on closing promoters. Next, we looked for regulators of the observed inflammatory response by performing motif discovery and enrichment using MEME-ChIP (**Fig. 5c; Source Data 9**). Opening OCRs were highly enriched for Spi1 binding sites. In our RNA-seq data, Spi1 was in the top 15% most expressed genes and upregulated in cGAS KO (LogFC = 0.4, p = 0.0145). Spi1 (Alias PU.1) is an ETS family transcription factor most known as a regulator of hematopoiesis. However, recent publications have demonstrated that Spi1 is expressed in fibroblasts in fibrotic diseases, and that it controls expression of multiple inflammasome genes^44,45^. Thus, the expression of inflammatory genes, as well as the fibrosis we observed in the liver in cGAS KO mice may be mediated by Spi1. Motif analysis on closing OCRs showed enrichment of motifs consistent with multiple forkhead box family transcription factors, primarily FoxE1 and FoxD2 (**Source Data 10**). FoxE1 is a factor involved in thyroid morphogenesis, whereas information on the role of FoxD2 is scarce. Thus, it is not clear to us how these factors may contribute to the cGAS KO phenotype.

**Figure 5.**
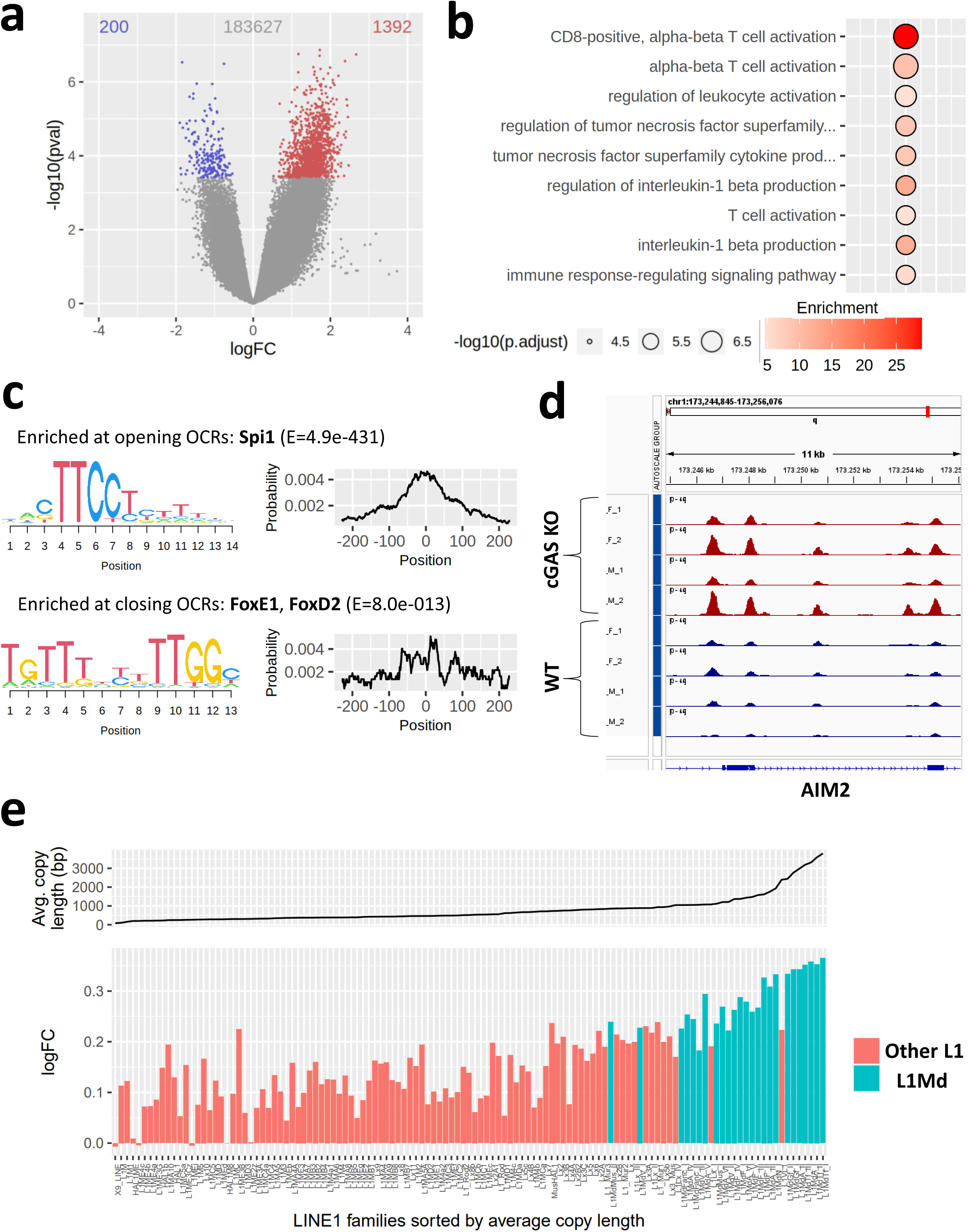
Primary cGAS knockout cells exhibit increased accessibility on pro-inflammatory genes and young L1s. (a) Volcano plot of differentially accessible OCRs in cGAS KO versus WT mouse primary lung cells. Relative to WT, cGAS KO samples display 1392 significantly more accessible OCRs and 200 less accessible OCRs. n = 4 mice for each group. (b) Top GO terms of differentially accessible promoters in cGAS KO versus WT mouse lung. (c) Motif discovery and enrichment using MEME-ChIP for regulators of the observed inflammatory response in cGAS KO cells. OCRs significantly more open in cGAS KO samples were enriched for Spi1 binding sites. OCRs significantly more closed in cGAS KO samples were most enriched at FoxE1 and FoxD2 binding sites. (d) AIM2 (one of the primary PYHIN genes; that stimulates expression of IL-18 and IL-6 via STING) is significantly more accessible in cGAS KO lung cells (red peaks) than in the WT lung cells (blue peaks). M/F indicate mouse sex. (e) Changes in accessibility of mouse LINE1 families upon cGAS KO. Families are sorted based on the average length of their copies, which is a proxy of evolutionary age. Teal bars indicate the evolutionary young L1Md clade. Y-axis is the fold change in cGAS KO ATACseq compared to WT (0 = no change).

In addition to increased pro-inflammatory gene accessibility, cGAS KO mice also displayed increased accessibility over L1 regions (**Fig. 5e**). The increase in accessibility of L1 elements in cGAS KO mice was highest for evolutionarily young L1 families, particularly in the L1Md clade. Because the youngest L1 families are more likely to be full-length and encode functional ORF1 and ORF2 proteins compared to older, truncated families, these elements are the most relevant drivers of L1 related pathologies, including inflammaging^7^. The increase in global chromatin accessibility on repetitive elements and particularly on younger L1s in cGAS KO cells suggests that nuclear cGAS is important for L1 repression.

### cGAS KO cells exhibit a global decrease in transposon DNA methylation

The striking changes in L1 gene expression/accessibility observed in RNAseq and ATACseq prompted us to investigate if cGAS KO mice display reduced genomic methylation compared to WT mice, as DNA methylation is one of the key mechanisms involved in transposable element repression. We utilized Nanopore sequencing to assess the global genomic changes in genomic methylation in cGAS KO and WT nuclei isolated from lung tissue.^46^ Specifically, we compared the average genome-wide methylation level for all transposable element families (**Fig. 6a**). Strikingly, cGAS KO mice exhibited a significant reduction in global methylation on most DNA, LINE, LTR, and SINE transposon families (**Fig. 6b**). Once again, multiple L1s within the L1Md clade showed significant loss of methylation, including the active L1MdA_I and L1MdV_I families (**Fig. 6c**). Intriguingly, almost all quantified SINE families showed significant demethylation (**Fig. 6d**). SINE elements encode no proteins, and instead mobilize by ‘hijacking’ the L1 ORF2p.^47,48,49^ Despite their inability to transpose autonomously, SINEs are among the most mobile TEs in mouse and human^50^. Therefore, the concurrent derepression of SINEs and LINEs is likely to cause severe cellular damage and may have contributed to the deleterious effects of cGAS KO.

**Figure 6.**
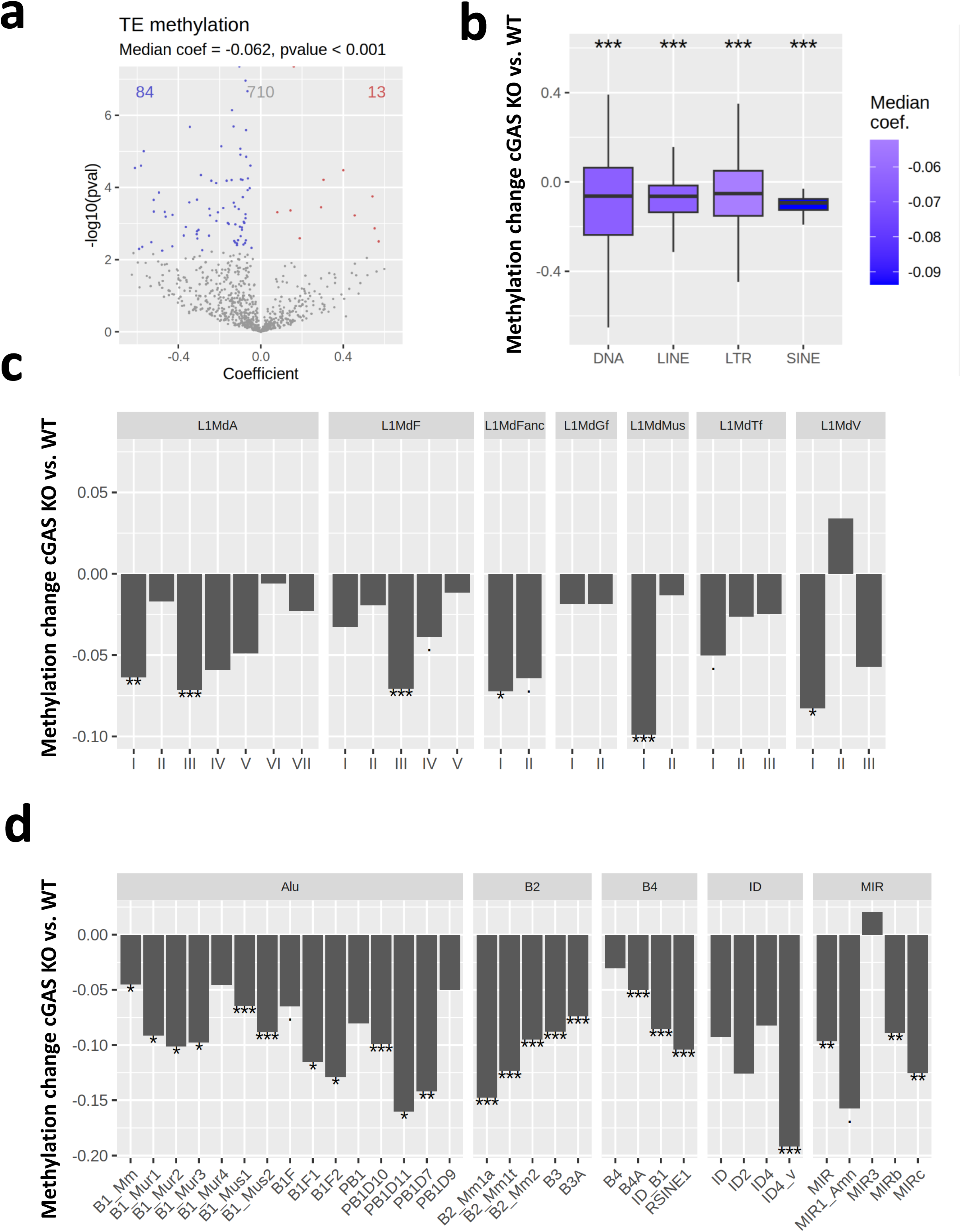
cGAS KO cells exhibit a decrease in methylation on genomic DNA. (a) Volcano plot of differentially methylated repetitive element families in cGAS KO versus WT primary mouse lung cells. Relative to WT, cGAS KO samples display 84 repeat families with significantly decreased DNA methylation. The majority of repeat families have lower methylation in cGAS KO lung tissue (One sample Wilcoxon signed-rank test). n = 3 mice for each genotype. (b) Change in genomic DNA methylation on various transposable element classes in cGAS KO relative to WT primary mouse lung cells. cGAS KO samples exhibit significant global reduction of DNA methylation on DNA, LINE, LTR, and SINE elements. Stars represent significance for one sample Wilcoxon signed-rank test (Null hypothesis: median logFC is zero). (c) Change in genomic DNA methylation on *Mus* L1Md elements in cGAS KO primary lung cells relative to WT. (d) Change in genomic DNA methylation on *Mus* SINE elements by family in cGAS KO primary lung cells relative to WT.

### Nuclear cGAS undergoes phase separation and is associated with heterochromatin

Given the increase in chromatin accessibility on repetitive elements along with the reduction of DNA methylation observed in the genome in cGAS KO cells, we hypothesized that nuclear cGAS might be involved in heterochromatin maintenance. Utilizing stimulated emission depletion (STED) microscopy, we visualized nuclear cGAS in primary mouse cells (**Fig. 7a**). Strikingly, cGAS formed condensed clusters in the nucleus, suggestive that liquid-liquid phase separation (LLPS) may stabilize cGAS-mediated genomic organization. WT human fibroblasts (HDFs) also displayed patterning indicative of LLPS for nuclear-localized cGAS (**Fig. 7b**). To verify that nuclear cGAS organization involves LLPS, we treated human fibroblasts with 1,6 Hexanediol (1,6 HD) to disrupt LLPS in cells (**Fig. 7b**). This treatment abolished the cGAS nuclear condensates, suggesting cGAS undergoes LLPS for proper localization on chromatin. Finally, we examined whether cGAS LLPS puncta co-localized with H3K9me3, a histone modification associated with heterochromatic, transcriptionally repressed regions of the genome^51^. The cGAS and H3K9me3 signals showed very strong co-localization (**Fig. 7a**). In contrast, cGAS KO cells lost H3K9me3 foci and displayed altered distribution H3K9me3 distribution pattern within the nucleus (**Fig. 7a, Sup. Fig. 6a**). This indicates cGAS is required for proper organization of H3K9me3 heterochromatin. Taken together, these data indicate cGAS colocalizes with heterochromatic histone marks and may be involved in heterochromatin maintenance.

**Figure 7.**
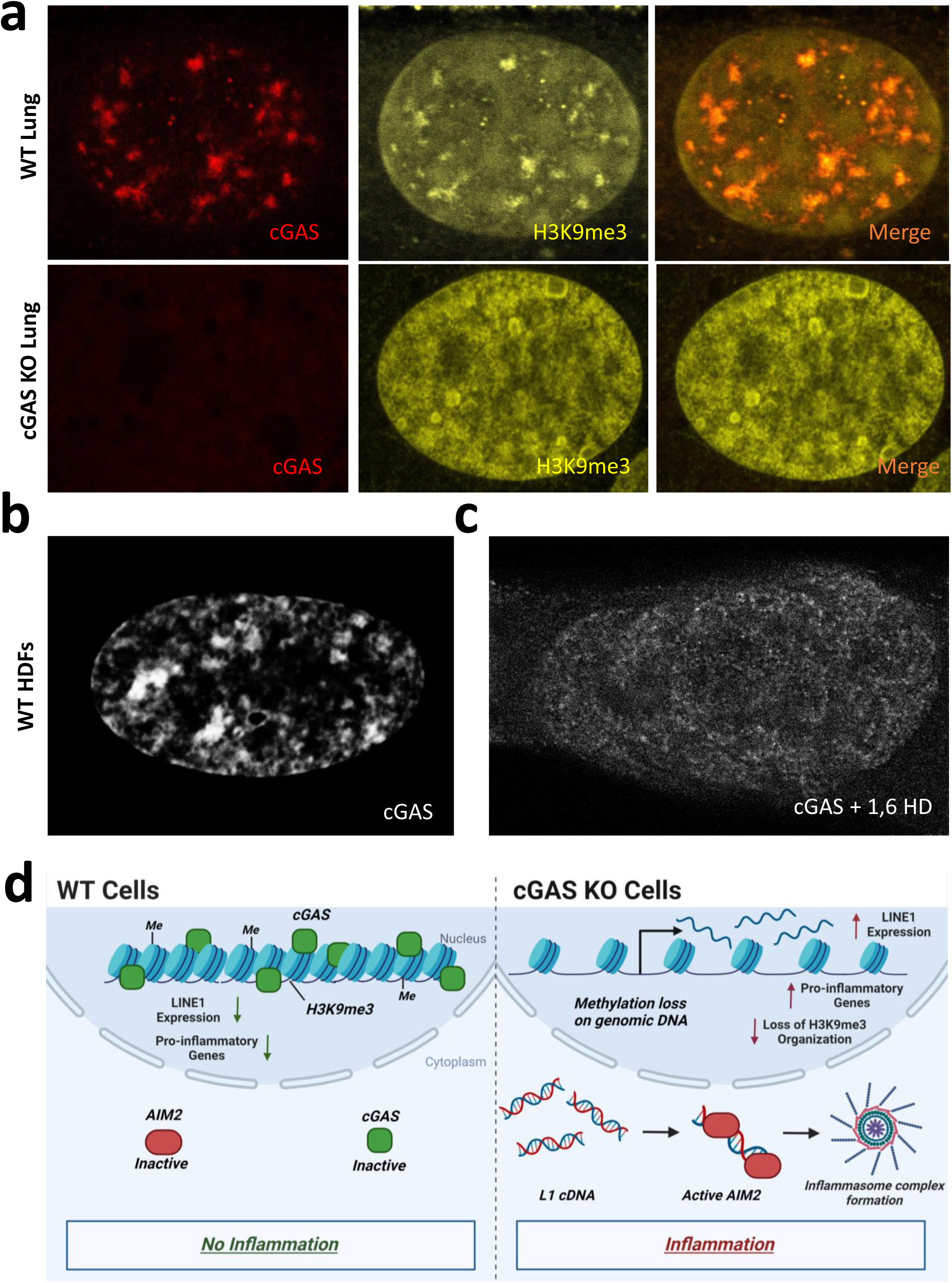
cGAS mediates chromatin organization. (a) Nuclear cGAS co-localizes with H3K9me3. STED images of primary lung cells from WT and cGAS KO mice stained for cGAS and DAPI. Experiments were performed in triplicate; n = 3 mice for each group. (b) cGAS forms LLPS condensates in the nucleus of human diploid fibroblasts. (c) 1,6 HD treatment disrupts cGAS LLPS nuclear condensates in human fibroblasts. (d) Model for nuclear cGAS function in maintaining chromatin organization and L1 repression. cGAS binding to nucleosomes promotes heterochromatin on L1 elements via phase separation. Loss of cGAS disrupts heterochromatin organization leading to L1 de-repression and induction of inflammation. Created in BioRender. Martinez, J. (2024) BioRender.com/x29g707

Loss of heterochromatin and de-repression of L1 elements associated with induction of inflammation represents an important hallmark of aging. Thus, our results suggest that cGAS KO mice display a progeroid phenotype. In summary, our results demonstrate a previously undescribed role for nuclear cGAS in maintaining heterochromatin and show that loss of cGAS leads to shortened healthspan associated with L1 derepression, DNA damage, and induction of inflammation (**Fig. 7f**).

## Discussion

Finding effective strategies to counteract age-related inflammation is important in order to improve healthspan and prevent age-related diseases. As cGAS/STING pathway is a key mediator of innate immune responses it is being actively explores as a target for pharmaceutical interventions^38,9,13,52^. However, the effects of cGAS depletion on longevity and long-term health have yet to be described. Our initial hypothesis was that cGas KO mice would display improved lifespan and healthspan. Unexpectedly, we found the opposite. Our data shows that cGAS deficient mice display premature aging phenotype characterized by increased frailty and massive induction of inflammation. Further analysis of cGAS KO mice revealed that their genomes undergo sweeping changes in chromatin organization that included more open chromatin, increase in overall transcription and de-repression of L1 elements. Importantly, we observed increased inflammation and L1 activation in both the cGAS KO mice and in cultured cells where cGAS was knocked down using siRNA, indicating that the observed inflammation is intrinsic to cGAS-deficient cells and is not the result of cGAS KO mice suffering from infections.

Since the demonstration that L1 elements are derepressed during aging and drive interferon response via the formation of cytoplasmic cDNAs, growing evidence links L1 elements to inflammation culminating in several clinical trials where L1 inhibition is probed and a therapy for age-related diseases^7,53,54^. It was recently demonstrated that L1s can undergo reverse transcription in the cytoplasm, hence elevated L1 expression leads to accumulation of L1 cDNAs which in turn lead to activated cGAS/STING pathway^38^. Surprisingly, we observed that cGAS KO cells accumulate high levels of L1 cDNAs and display elevated innate immune response. We hypothesize that other DNA sensors compensate for the absence of cGAS. Indeed, we observed elevated levels of Aim2 gene expression in cGAS KO cells.

Why is the loss of cGAS associated with L1 derepression? Interestingly, we did not observe L1 activation in STING or PYHYN in KO mice. While the best understood role of cGAS is in the cytoplasm, a larger fraction of cGAS is localized to the nucleus^42^. Nuclear cGAS binds to chromatin where it does not induce interferon response as this function is repressed by binding to nucleosomes^55^. Several roles were proposed for nuclear cGAS which include repressing DNA repair, slowing down transcription forks, and repressing L1 elements via targeted degradation of ORF2p^43,56,57^. Here we show that cGAS co-localizes with heterochromatin marks and loss of cGAS leads to changes in the distribution of these marks and generalized opening of chromatin and L1 derepression. We propose that cGAS plays a role in maintaining chromatin organization and repressing repetitive elements. This function is consistent with previous observations where cGAS was shown to repress homologous recombination and slowdown of transcription forks, as generalized opening of chromatin would both allow for more active recombination and increase the speed of transcription. We also show evidence that cGAS undergoes LLPS in the nucleus. It was previously shown that cytoplasmic cGAS undergoes LLPS upon binding to DNA^58^. Importantly, the nucleosome-binding domain, rather than the cGAMP producing domain is required for phase separation of cGAS^58^. While LLPS with naked DNA triggers immune signaling, it is possible that LLPS with nucleosome bound DNA is immunologically silent and promotes compaction of heterochromatin.

Intriguingly, differences across the most important residues in nucleosome binding domain across animal species raises the possibility that the nucleosome binding efficiency of cGAS may play a role in longevity by promoting chromatin maintenance and reducing L1 activity and chronic inflammation. Many species of bats have dramatically longer lifespans than could be expected for an organism of their size^59^. One of the hypotheses for this extended longevity in bats is a reduction in inflammaging due to better control of harmful immune signaling. Bats have downregulated multiple components of DNA sensing pathways including complete loss of PYHIN family of cytosolic sensors^60^. Additionally, bat STING has lost the residue S358, which has been shown to be important in amplifying IFN response following residue phosphorylation upon STING activation^61^. In defiance of this trend, bats have not only retained their cGAS, but also contain intriguing differences in key residues involved in nucleosome-binding that may alter the affinity of bat cGAS in nucleosome binding. Further exploration of cross-species cGAS functionality and connections to longevity are warranted in future studies.

Collectively, this study has established a previously undescribed role for nuclear cGAS in maintaining heterochromatin organization via LLPS. We show that loss of nuclear cGAS results in increases in L1 mRNA transcription and the subsequent cDNA and ORF1p accumulation in the cytoplasm. Loss of nuclear cGAS also leads to increased genomic accessibility of pro-inflammatory genes. As mitigation of dysregulated inflammatory activity continues to be pursued as a therapeutic intervention, the effect of potential treatments on the stability and maintenance of nuclear cGAS must be approached with caution.

It is tempting to speculate that original evolutionary function of cGAS was to suppress transposable elements. Hence, most cGAS is localized in the nucleus, and when transposons escape into cytoplasm cGAS follows them. A complete loss of cGAS causes reactivation of these genomic parasites, which cannot be compensated by other silencing factors.

## Materials and Methods

### Animal Care

cGAS KO, STING KO, and wild-type mice were obtained from Jackson Laboratories (strain B6-cGas^tm1d(EUCOMM)Hmgu^/J, #026554 and strain B6-Sting^tm1.2Camb^/J, #025805) and cared for in accordance with the regulations developed and approved by the University of Rochester Committee on Animal Resources (UCAR). UCAR is compliant with FDA and NIH (National Institutes of Health) animal care guidelines and reviews all protocols before approval and implementation (#2017-027). Animals were housed in a one-way facility utilizing microisolator technology. Frailty scoring conducted according to established protocol^30^.

### Tissue histology, hematoxylin & eosin (H&E) staining, and imaging

Mice were sacrificed via isoflurane exposure and perfused with PBS. Tissues were then isolated from mice, placed in a histology cassette, and fixed in 4% paraformaldehyde (PFA) for 48 hours. Tissues were subsequently submitted to the University of Rochester CMSR Histology Core (NIH P30 AR069655) for paraffin embedment, sectioning, and H&E staining. Slides were imaged using brightfield microscopy.

### Tissue processing and RNA extraction

Tissues were isolated from mice, flash frozen in liquid nitrogen, then pulverized using piton/cell crusher chilled using liquid nitrogen to a fine powder. RNA was isolated from pulverized tissue using Trizol. Insoluble material was removed by spinning tubes for 1 minute at 12,000 g/4-degrees and transferring supernatant to new tube. 100 µl chloroform was added to this solution, shaken vigorously for 15 seconds, incubated at room temperature for 5 minutes, then centrifuged for 15 minutes at 12,000 g. Aqueous phase containing RNA was then extracted to a new tube, mixed with 250 µl isopropanol, then incubated at room temperature for 10 minutes. The samples were then centrifuged for 10 minutes at 12,000 g. Supernatant was then removed, leaving behind RNA pellet. The pellet was then washed in 500 µl 75% ethanol in DPEC ddH2O and centrifuged at 8,000 g for 5 minutes. EtOH was then removed, and the pellet allowed to air dry for 10 minutes. RNA was further purified utilizing RNA PureLink Mini Kit (Thermo). cDNA was then generated using Superscript III (Thermo-Fisher) cDNA kit using random hexamer primers. qRT-PCR was performed using the Bio-Rad CFX Connect Real Time machine and SYBR Green Master Mix (Bio-Rad) using 30 ng/cDNA per reaction.

### Cell Culture

All cell lines were grown in humidified incubators at 5% CO2 and 5% O2 at 37 degrees Celsius. Mouse embryonic fibroblasts (MEFs) were cultured in Eagle’s minimum essential medium with 15% fetal bovine serum (FBS) and 1X penicillin/streptomycin. Primary lung cells were cultured in Dulbecco’s modified Eagle’s medium nutrient mixture F12 with 15% FBS and 1X antibiotic-antimycotic. Cell lines are regularly tested for mycoplasma contamination.

### cGAS siRNA treatment

cGAS siRNA (Thermo-Fisher) was delivered to MEFs utilizing Lipofectamine RNAiMAX Transfection Reagent (Invitrogen) following manufacturer protocol. Cells were fixed and stained (either via immunofluorescence or DNA smiFISH) 72-hours post transfection. cGAS knockdown was verified via western blot and qRT-PCR.

### Quantitative RT-PCR from cells

Total RNA from cultured cells was isolated using RNeasy kit from Qiagen following kit protocol. cDNA was then generated using Superscript III (Thermo-Fisher) cDNA kit using random hexamer primers. qRT-PCR was performed using the Bio-Rad CFX Connect Real Time machine and SYBR Green Master Mix (Bio-Rad) using 30 ng/cDNA per reaction. Each sample was run in triplicates. Mouse L1 primers targeted towards evolutionarily young L1MdA elements previously described, developed, and optimized were used for L1 expression comparison and standardized to QuantumRNA Actin Universal primers^7,24^. Primers utilized: mL1 Fwd-ATGGCGAAAGGCAAACGTAAG; mL1 Rev-ATTTTCGGTTGTGTTGGGGTG; 5S Fwd-CTCGTCTGATCTCGGAAGCTAAG; 5S Rev-GCGGTCTCCCATCCAAGTAC.

### Immunofluorescence Staining and Imaging

24 hours before fixation, 40,000 cells were seeded per chamber in a 2 well chamber slide (Thermo-Scientific Nunc Lab-Tek II Chamber Slide System, 2 well) and cultured under previously described cell culture conditions. Cells were washed twice with PBS before fixing with 3.7% PFA and placed in 37-degree incubator (5% O2 and 5% CO2) for 10 minutes. After 3 washes with PBS, cells were permeabilized using 0.5% Triton X-100 (in PBS) while shaking at room temperature for 10 minutes. Cells were then washed 4 times with PBS for 5 minutes each. If cells were to be stained using the antibody targeting DNA-RNA hybrids, slides were wrapped in parafilm and further fixated in 100% methanol overnight at -20-degrees. Cells were then incubated for one hour at room temperature using blocking buffer (10% FBS and 5% BSA in PBS and filter sterilized). Appropriate primary antibodies were added and incubated with shaking at 4-degrees overnight: cGAS (1:1,000, cell signaling), γH2AX (1:500, ABCAM), or ORF1p (1:100, Boeke Lab). After washing cells with 0.1% Triton X-100 4 times at room temperature for 5 minutes each, secondary antibodies were added for one hour at room temperature for either Alexa-488 or Alexa-568 (1:1,000, Invitrogen). Cells were further washed 4 times with PBS for 5 minutes each before mounting using DAPI mounting medium (Sigma-Aldrich). After drying, slides were imaged using confocal microscope.

### DNA smiFISH

Protocol for DNA smiFISH was adapted from original protocol^62^. 2 well chamber slides were utilized in leu of coverslips. Cells were seeded one day prior to fixation for 75% confluency. For fixation, media was removed, and cells washed twice with PBS before fixing in 4% paraformaldehyde for 10 minutes at 37-degrees. Cells were then permeabilized in 70% ethanol overnight at 4-degrees. Cells were washed with PBS then incubated in 15% formamide in 1X SSC for 15 minutes at room temperature. 50 µl of hybridization buffer (comprised of 20X SSC, 20 µg/µl yeast tRNA, 100% formamide, fluorescently labeled probes, 20 mg/mL ultra-pure BSA, 200 mM VRC, 40% dextran sulphate, H2O) was placed on a petri dish and slides were placed onto the dish facing down. Cells were incubated at 37-degrees overnight with a 3.5 cm petri dish containing 15% formamide in 1X SSC inside the petri dish containing slides to prevent dehydration. The next day, cells were washed twice for 30 minutes at 37-degrees in 15% formamide/1X SSC. Prior to mounting, cells were washed twice in PBS before 35 µl/chamber of Vectashield mounting medium with DAPI were applied to cells. Slides were stored at -20-degrees long-term.

### Protein Isolation and Western Blotting

Protein from cultured cells were isolated utilizing RIPA lysis and extraction buffer (Thermo-Scientific) in the presence of protease and phosphatase inhibitors (Sigma-Aldrich). After resuspension in RIPA, cells were incubated with rotation at 4-degrees for 30 minutes and then centrifuged at maximum-speed in a microcentrifuge for 15 minutes at 4-degrees. Isolated proteins were collected via supernatant and stored at -20-degrees until use. Supernatant was mixed 1:1 with 2X Laemmli buffer and boiled for 5 minutes at 95-degrees before being centrifuged for one minute at 14,000 RPM and loaded into a Bio-Rad 4%-20% gel. Proteins were then transferred to a PDVF membrane and blocked with Intercept TBS Blocking Buffer for 1 hour at room temperature. Membranes were incubated overnight at 4-degrees with shaking using appropriate antibodies: cGas (1:1,000, cell signaling), vinculin (1:1,000, cell signaling), H3 (1:10,000, Abcam), or ORF1p (1:500, Boeke Lab). Membranes were washed with TBST 3 times for 5 minutes each before applying secondary fluorescent antibody (1:2,500, StarBright Blue 520 Bio-Rad) for 1 hour at room temperature. Membranes were then washed and imaged using Bio-Rad ChemiDoc.

### Sub-Cellular Protein Fractionation

Subcellular protein fractionation for cultured cells was performed according to protocol provided by Thermo-Scientific. The soluble nuclear and chromatin bound protein fractions were combined to comprise the ‘nuclear’ fraction. Fractionation efficiency was verified by western blot.

### STED Staining and Imaging

Cells were plated on STED compatible, glass bottom 35cm cell culture dishes and grown to 80% confluency. Media was removed and cells washed twice with PBS. Cells were fixed in 4% paraformaldehyde at 37-degrees for 10 minutes and then washed twice in PBS. Cells were permeabilized in 0.5% Triton X-100 at room temperature for 10 minutes then washed 3 times in PBS then once more for 5 minutes. Cells were then blocked (10% FBS, 5% BSA in PBS and filter sterilized) for one hour. 100 µl of primary antibody solution (antibodies diluted 1:100 in blocking buffer) was applied directly to glass portion of dishes and incubated overnight at 4-degrees with shaking. The next day, primary antibody was removed, and cells washed 4 times in 0.1% Triton X-100 at room temperature for 5 minutes. 100 µl of secondary antibody solution (antibodies diluted 1:100 in blocking buffer) was applied directly to glass and shaken at room temperature for 1 hour (covered in dark box). Cells were then washed twice with 0.1% Triton X-100 then twice in PBS for 5 minutes. DAPI was applied to cells by diluting DAPI 1:1000 in PBS and shaken at room temperature for 15 minutes. Finally, cells were washed twice in PBS before applying 3 drops (∼75 µl) Prolong Gold antifade to the glass and apply coverslip. Dishes were stored at 4-degrees in the dark until imaging. Cells were imaged using the Abberior easy3D STED microscope.

### RNA-seq analysis

Samples were sequenced at an average depth of 50 M paired end reads. Read trimming and adapter removal was performed with TrimGalore! Trimmed reads were aligned to the GRCm39 mouse genome using STAR^63^ with the following arguments: --outFilterMultimapNmax 100--winAnchorMultimapNmax 200 --outFilterMismatchNoverLmax 0.04 to retain multimapping reads as recommended by TEtranscripts^64^. Gene and TE expression was then quantified using TEcounts (part of TE transcripts) using the curated GTF file available for GRCm39 on the TEtranscripts website. The remaining analyses were performed in R 4.2.2. Differential expression was evaluated using EdgeR^65^ using the model ∼Genotype+Sex and GSEA^66^ was performed with ClusterProfiler^67^.

### ATAC-seq analysis

Samples were sequenced at an average depth of 50 M paired end reads. Read trimming and adapter removal was performed with TrimGalore! Trimmed reads were aligned to the GRCm39 mouse genome using Bowtie2^68^ with arguments --very-sensitive -X 1000 –dovetail. Only proper pairs and primary alignments were retained using samtools^69^ view -f 0x2 -F 0x100. This has the effect of retaining only one random alignment among multimapping reads, as recommended by Teissandier et al^70^. Mitochondrial reads and PCR duplicates were removed using samtools and picard. Cleaned bam files were converted to bed format and used to call peaks with MACS2 with arguments -f BED --keep-dup “all” -q 0.01 --nomodel --shift -100 --extsize 200 for each sample individually. A unified peak set was generated as previously shown^71^ by first merging the peaksets of all samples using bedtools^72^ merge. Next, we identified regions which were called as peak in at least 2 samples using bedtools multiinter. Finally, we only kept peaks within the union peakset if they contained at least one of the regions called in 2 samples or more using bedtools intersect. Repetitive element annotation was performed with RepeatMasker. Accessibility of OCRs and repetitive elements was simultaneously quantified using featureCounts^73^ in the subread package, with arguments -F SAF -p -B --read2pos 5 -O -- fraction, which reduces reads to their 5’ end (Tn5 cut site) and counts reads fractionally, if they overlap both a peak region and a repeat region. Differential accessibility was evaluated using EdgeR^65^ using the model ∼Genotype+Sex. Annotation of OCRs was performed with ChipSeeker^74^. BigWig coverage tracks were generated using deeptools^75^ with scale factors calculated as (1e6 / #reads within peaks). Motif analysis was performed with MEME-ChIP^63^.

### Nanopore methylation analysis

DNA samples were prepared using the Oxford Nanopore rapid barcoding kit RBK114-24 following the manufacturer’s protocol and sequenced on a MinION Mk1B. Basecalling was performed using dorado version 0.7.3, using the dna_r10.4.1_e8.2_400bps_sup@v5.0.0 basecalling model, calling 5mC and 5hmC in a CpG context (5mCG_5hmCG). Reads were aligned to GRCm39 using dorado aligner, demultiplexed with dorado demux --no-classify --no- trim. Unmapped reads, secondary and supplementary alignments were removed using samtools^69^ view -F 0x904 and CpG methylation was quantified using modbam2bed --extended - -cpg --aggregate. Finally, we grouped CpGs based on the type of repetitive element they were contained within (RepeatMasker annotation), and only considered repetitive element families quantified by an average of 50 or more CpGs across samples. Differential methylation was evaluated using a binomial generalized linear model fitted to the count of methylated and unmethylated reads for each repetitive element family (cbind(metCount, unmetCount) ∼ Genotype).

### Quantification and Statistical Analysis

Unless specified otherwise, the student’s t-test was utilized for comparison between groups. All tests were two-tailed, and p-values were considered significant below the 0.05 threshold. Multiple testing corrections were performed with the Benjamini-Hochberg procedure when applicable.

### Code Availability

The code used for preprocessing and analyzing rna-seq, atac-seq and nanopore data is available at https://github.com/SunScript0/cGASKO-inflam.

## Author Information

### Author Contributions

JM, FM, AS, and VG conceived of the experiments. JM, MM, LF, NS, SJB, EH, JC, VP, and SAB performed the experiments. JM, FM, AS, and VG analyzed data and FM conducted bioinformatic analyses. JM, FM, AS, and VG wrote the manuscript with input from all authors.

### Data accessibility

Data will be made available upon manuscript publication.

## Acknowledgements

The authors would like to thank all members of the Gorbunova and Seluanov laboratories for helpful discussions. We would also like to thank the Sedivy lab for their support and contributions to this work and the Boeke lab for their lending of the LINE1 ORF1p antibody (EA13).

