## Supplemental Figures for "cGAS deficient mice display premature aging associated with de-repression of LINE1 elements and inflammation"

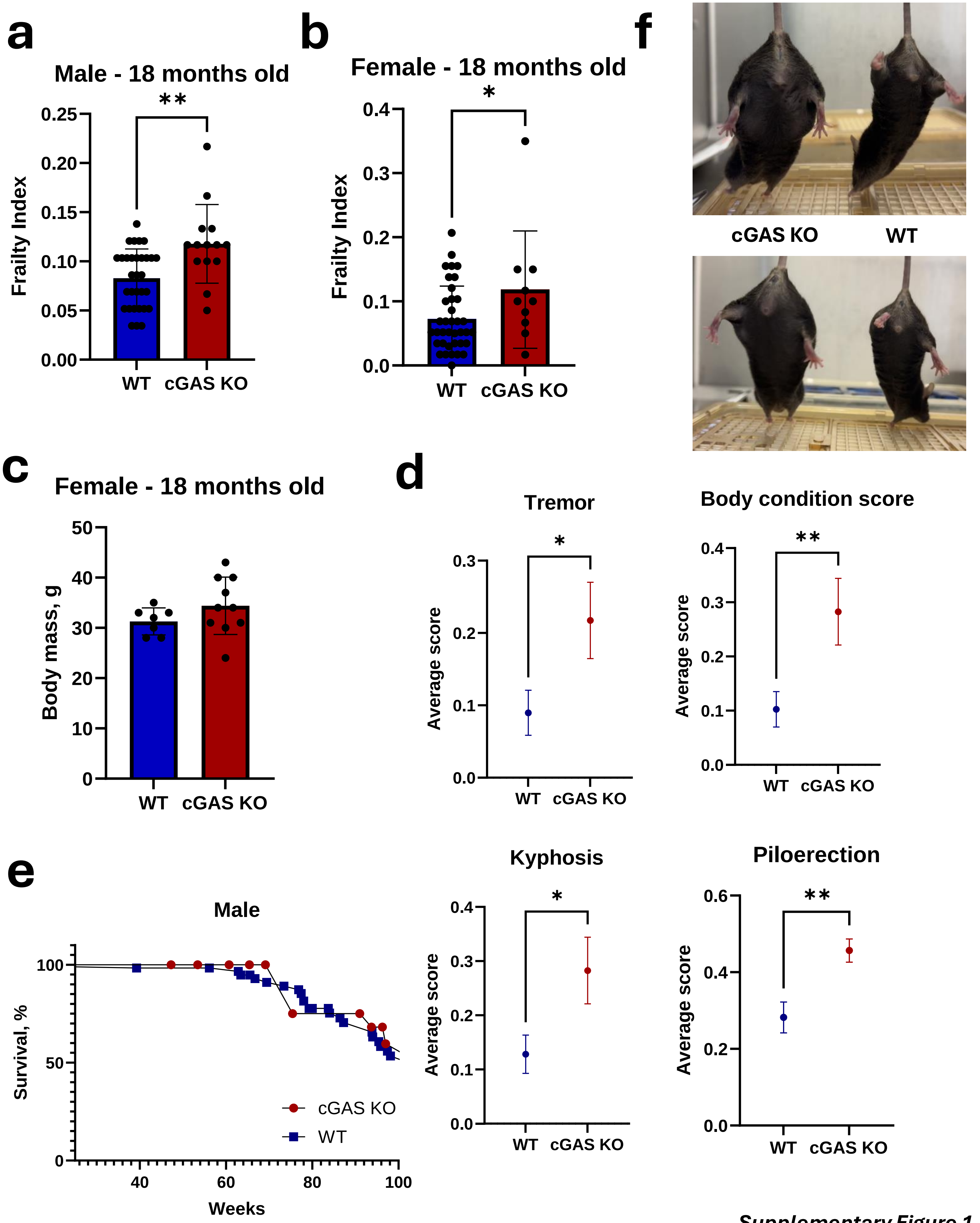


*Supplementary Figure 1*

**Supplementary Figure 1. cGAS KO mice display increased frailty.**

cGAS KO males and females display higher frailty score with age. *p < 0.05, **p < 0.01.

(a) Frailty index of 18-months-old cGAS KO and WT male mice. WT n = 31, cGAS KO n =14.
(b) Frailty index of 18-months-old cGAS KO and WT female mice WT n = 37, cGAS KO female n = 10.
(c) Body mass of female cGAS KO mice. WT n = 7, cGAS KO n = 10.
(d) Individual components of the frailty index where cGAS KO mice display worse scores. Kyphosis, piloerection, body condition and tremors at 18-months of age. Graphs display average score for each parameter along with standard error of the mean (SEM). *p < 0.05, **p < 0.01, t-test.
(e) Survival of WT and cGAS KO male mice. (cGAS KO n = 44 mice, WT n = 60 mice; WT median survival = 101.7 weeks; cGAS KO median survival = 97.7 weeks; p = 0.32, Mantel-Cox test).
(f) Representative images of cGAS KO and WT male mice at 14-months-old. cGAS KO mouse weight is 52.69g; WT mouse weight is 31.98g (left pair). cGAS KO mouse wight is 52.61g; WT mouse weight is 34.22g (right pair).

a

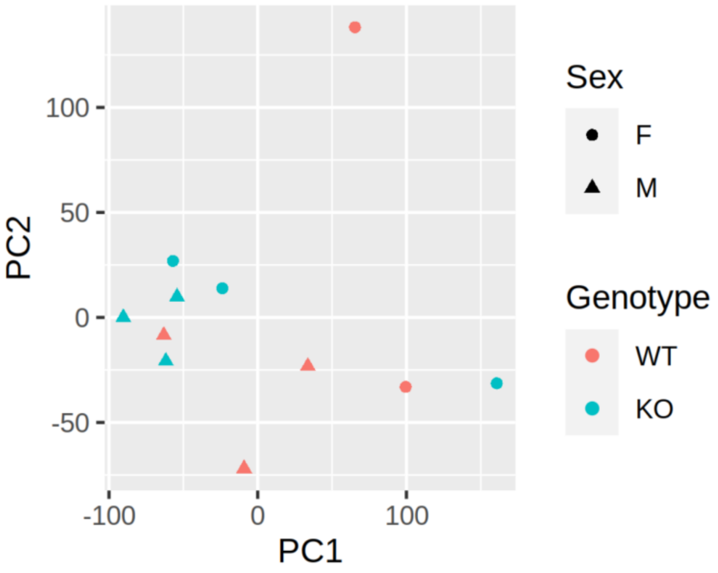


b

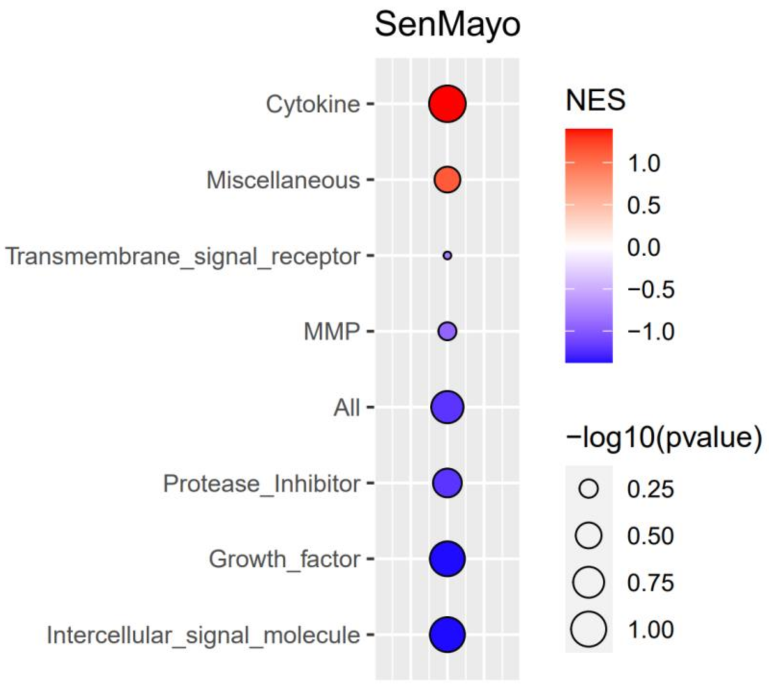


**Supplementary Figure 2. RNAseq reveals pro-inflammatory signature in cGAS KO lung.**

(a) PCA analysis of RNAseq samples of WT and cGAS KO samples colored by genotype and sex. cGAS KO n = 6 mice; WT n = 5 mice.

(b) SenMayo senescence panel for cGAS KO vs WT samples.

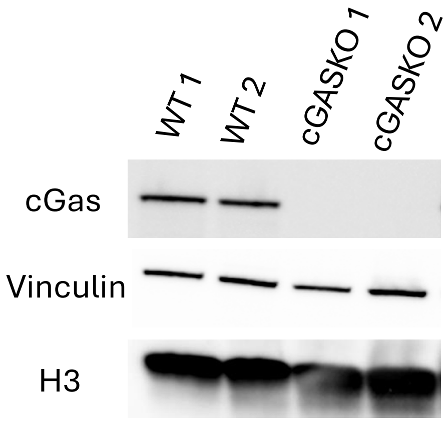


**Supplementary Figure 3. Western blotting confirms cGAS KO.**

Western blot analysis of cGAS KO and WT lung tissue. H3 and vinculin were used as a loading controls.

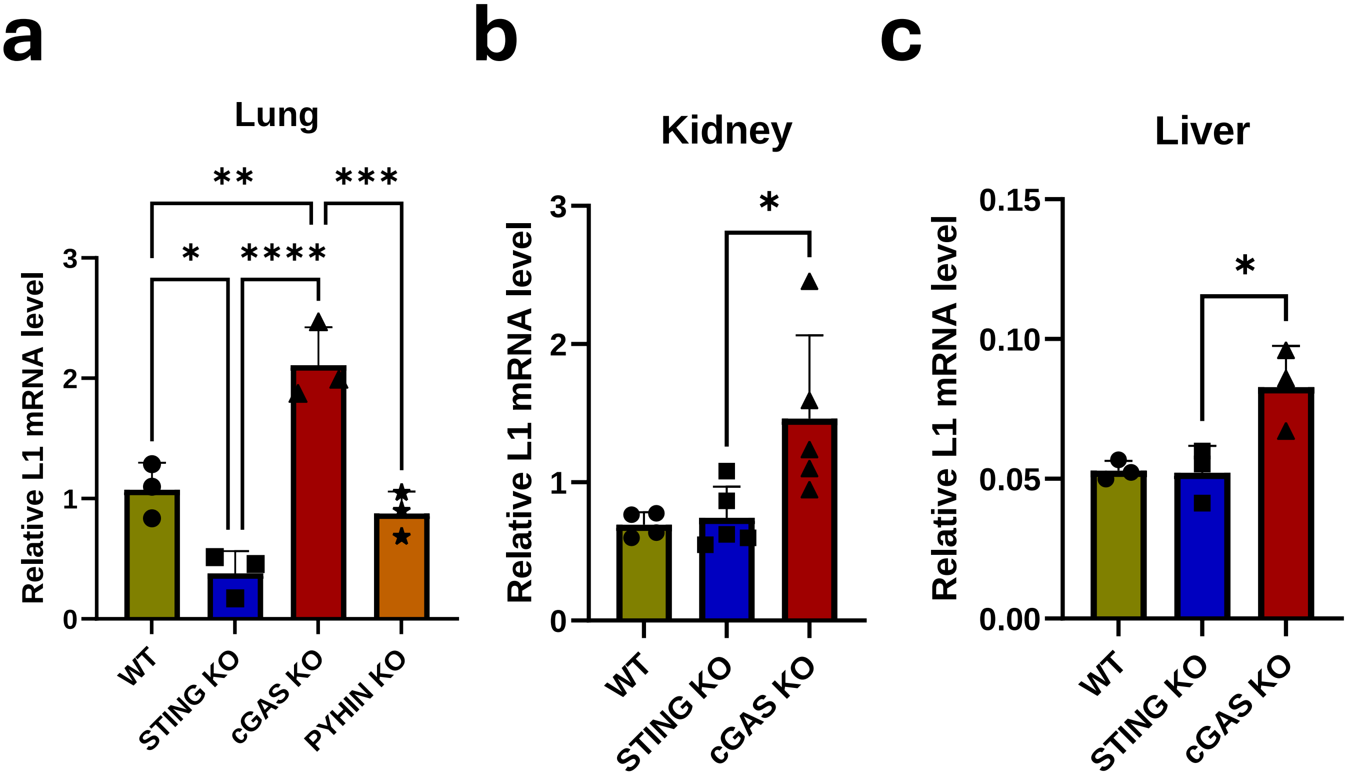


**Supplementary Figure 4. Depletion of other cytoplasmic DNA sensors does not lead to L1 activation.**

L1 mRNA is elevated in several tissues in cGAS KO mice but not PYHIN KO or STING KO. Organs were harvested from 12-month-old WT and cGAS KO mice. L1 expression was measured via RT-qPCR and normalized to EEF2.

(a) Lung. *$p < 0.05$, **$p < 0.01$, ***$p < 0.001$, ****$p < 0.0001$, t-test.

(b) Kidney. *$p < 0.05$, t-test.

(c) Liver. *$p < 0.05$, t-test.

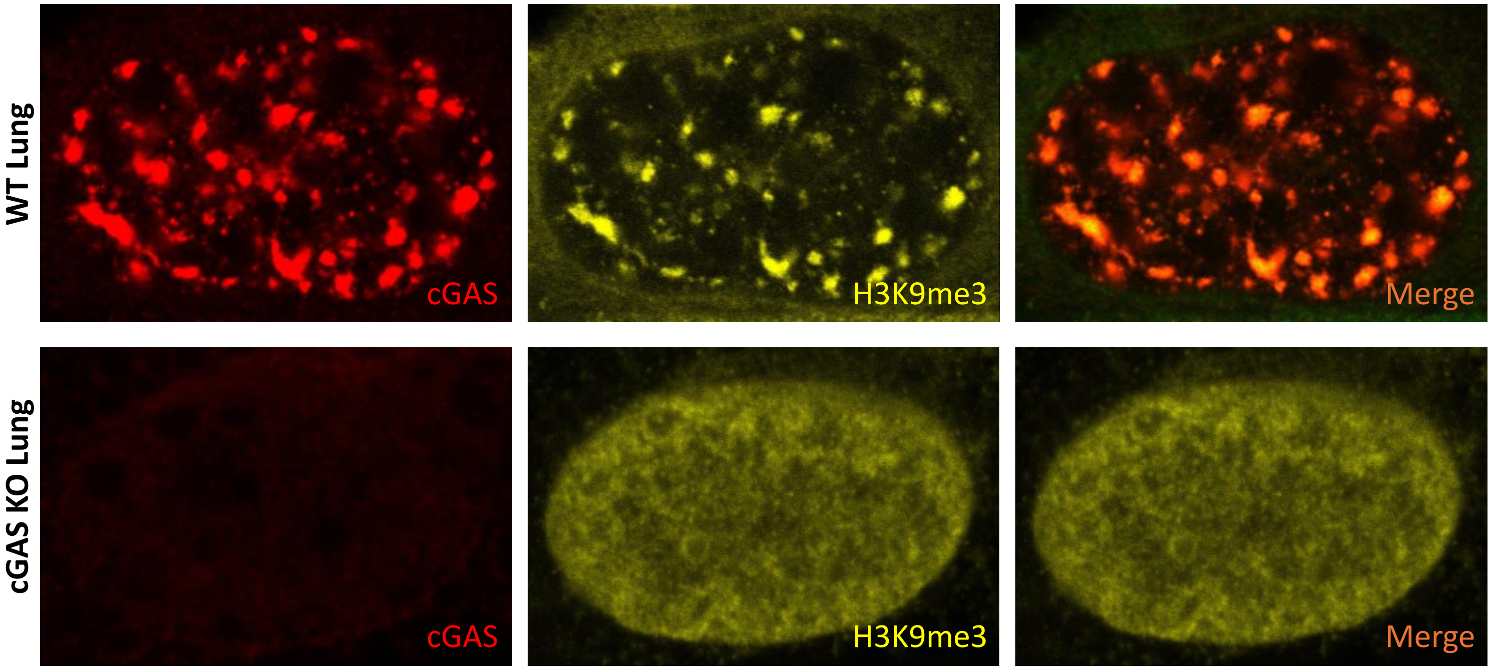


**Supplementary Figure 5. Loss of cGAS results in H3K9me3 disorganization in the nucleus.**

cGAS KO primary lung fibroblasts were stained with antibodies to H3K9me3 mark of heterochromatin. cGAS KO cells display aberrant distribution of H3K9me3 with the loss of heterochromatin clusters. Cells were isolated from 12-month-old male WT and cGAS KO mice.
